# *Sanctuary*: A *Starship* transposon facilitating the movement of the virulence factor ToxA in fungal wheat pathogens

**DOI:** 10.1101/2024.03.04.583430

**Authors:** Angus Bucknell, Hannah M. Wilson, Karen C. Gonçalves do Santos, Steven Simpfendorfer, Andrew Milgate, Hugo Germain, Peter S. Solomon, Adam Bentham, Megan C. McDonald

## Abstract

There is increasing evidence that mobile genetic elements can drive the emergence of pathogenic fungal species by moving virulence genes horizontally. The 14 kbp *ToxhAT* transposon has been shown to be moving the necrotrophic effector, *ToxA,* horizontally between fungal species that infect *Triticum aestivum* (wheat), namely *Parastagonospora nodorum*, *Pyrenophora tritici-repentis*, and *Bipolaris sorokiniana*. All three species utilise the ToxA protein to infect wheat. Previous genomic evidence found *ToxhAT* in distinct chromosomal positions in two isolates of *B. sorokiniana*, indicating that the transposon is still active in this species. Here we confirm the movement of *ToxhAT* using long-read Nanopore MinION sequencing of eight novel and one previously published *B. sorokiniana* isolates. One event of independent transposition of *ToxhAT* was observed, and target site duplications of “TA” were identified, confirming this was an autonomous movement facilitated by a yet unidentified transposase. Whole genome analysis revealed that *ToxhAT* is a passenger embedded in a much larger, conserved 170–196 kbp mobile genetic element. This element, termed *Sanctuary*, belongs to the newly described *Starship* transposon superfamily. This classification is based on the presence of short direct repeats, empty insertion sites, a putative tyrosine recombinase gene and other features of *Starship* transposons. We also show that *ToxhAT* has been independently captured by two different *Starships*, *Sanctuary* and *Horizon* which share little to no sequenced identity, outside of *ToxhAT.* This classification makes *Horizon* and *Sanctuary* part of a growing number of *Starships* involved in the horizontal gene transfer of adaptive genetic material between fungal species.

**Importance:** The work presented here expands our understanding of a novel group of mobile genetic elements called *Starships* that facilitate the horizontal exchange of virulence genes in fungal pathogens. Our analysis shows that *Sanctuary* and *ToxhAT* are likely active and autonomous transposons in the *B. sorokiniana* genome. We also show that the smaller *ToxhAT* transposon has been independently captured by two different *Starships*, *viz. Sanctuary* in *B. sorokiniana* and *Horizon* in *P. tritici-repentis* and *P. nodorum.* Outside of *ToxhAT* these two *Starships* share no sequence identity. The capture of *ToxhAT* by two different mobile elements in three different fungal wheat pathogens demonstrates how horizontal transposon transfer is driving the evolution of virulence in these important wheat pathogens.

## Introduction

Horizontal gene transfer (HGT) is the non-Mendelian exchange of genetic material between organisms (1, 2). HGT is a significant driver of rapid adaptation to a changing environment and is of particular concern in microbial pathogens (3). Gene exchange through HGT can rapidly enhance virulence (4), increase host range (5), or increase microbial competitive ability (6). Eukaryotic HGT was once thought to be a rare event due to the complexity of these cell types (7). However, as more genome assemblies became available, especially ones assembled with long-read sequencing technologies, many examples of HGT between eukaryotes have been described (1). Long-read assemblies have also enabled a much more detailed look at the repetitive content of genomes, namely transposons and their history of HGT. While such discoveries were made before the advent of next-generation sequencing, dating back to the 1990s, robust evidence in the form of chromosome-level genome assemblies has uncovered many previously overlooked HGT events (8). These discoveries suggest that transposon-mediated HGT between eukaryotes is an ongoing and evolutionarily ancient form of adaptive evolution, where the underlying mechanism of how these transposons move between organisms remains to be characterised (1, 2).

As opposed to movement between different species, transposon movement within a genome is fairly well understood. Transposons are broadly classified into two major classes, based on whether they use an RNA (Class I) or DNA (Class II) intermediate to move locations within a genome (9). Other characteristics, such as conserved encoded proteins, structural motifs and target site preferences are used to further divide these transposons into orders, superfamilies and families (9). Class I retrotransposons are characterised by transposition involving an RNA intermediate. These transposons are often flanked by Long-terminal repeats (LTR), that facilitate their movement. Class II DNA transposons have no RNA intermediate and move through excision and integration. Class II transposition is facilitated by a transposase or recombinase, and are often bound by terminal inverted repeats (TIR) (9, 10). These TIRs define transposon boundaries and contain the recognition sites for the transposase that cleaves the DNA (9, 11). Target site duplications (TSD) are another important non-coding feature of both Class I and II transposons. The length and composition of LTR, TIR, and TSD regions are specific to the enzymes facilitating movement, are conserved within transposon families, and are used to classify novel transposons (9, 12).

Insertion of a transposon into a new genomic location usually involves a double-stranded break of the DNA that creates short overhangs. The backfilling of these overhangs leads to a duplication of this sequence on either side of the new insertion site. In many instances these TSDs can be quite difficult to recognise as they range in size from 2-8 base pairs. One useful technique to help identify TSDs is to compare the DNA sequence flanking the insertion to another individual that does not carry the transposon at the same site, often referred to as “empty-sites” (10, 13). This sequence-based evidence is only a snapshot of the transposon in the genome at a given time, however in the absence of an experimental system where all transposon locations can be precisely determined at different life stages of an organism, these TSDs remain the best way to identify active transposon families within a genome.

One well studied example of eukaryotic HGT is the *ToxA* virulence gene. *ToxA* is found in the fungal wheat pathogens *Pyrenophora tritici-repentis* (14), *Parastagonospora nodorum* (15), and *Bipolaris sorokiniana* (16), all of which cause severe crop losses in wheat crops globally (17, 18). These pathogens can all infect wheat without *ToxA,* however presence of this effector has been correlated with more severe disease symptoms indicating this gene can confer a strong fitness advantage (19). Long-read sequencing was used to demonstrate that *ToxA* is carried within a conserved transposon that was horizontally transferred between *P. nodorum* and *P. tritici-repentis*, and *B. sorokiniana* (20). This transposon, now called *ToxhAT*, is a single-copy, 14 kbp Class II transposon that is found in all three fungal species. *ToxhAT* remains highly conserved with a sequence similarity of ∼92% at the nucleotide level, indicative of a very recent HGT event (16). Homologous DNA upstream (61.2 kbp) and downstream (1.7 kbp) of *ToxhAT* is shared between *P. tritici-repentis* and *P. nodorum*, suggesting the HGT event was greater than just *ToxhAT*. However, between *B. sorokiniana* and these other two species, only *ToxhAT* is shared (20). Questions such as the order of acquisition by each species, and the origins of *ToxhAT* and *ToxA* remain (21).

*ToxhAT* in *P. tritici-repentis* and *P. nodorum* is believed to be inactive due to a high level of repeat induced point (RIP) mutation that disrupt the TIRs of *ToxhAT* in these species (20). RIP is a mechanism of fungal genome defence where cytosine to thymine (C to T) and guanine to adenine (G to A) polymorphisms are induced (22). Minimal RIP mutation of *ToxhAT* in *B. sorokiniana* has left the transposon largely intact and recent data from one genome assembly has indicated that *ToxhAT* remains active (20). In this *B. sorokiniana* isolate, the exact 14 kbp region of *ToxhAT*, bounded by the TIRs, was inverted and located on a different chromosome in reference to the other two isolates, indicative of active transposition. In another isolate, a roughly 200 kbp region of DNA including *ToxhAT* was found translocated to another chromosome, indicative of chromosomal translocation or the movement of a larger mobile genetic element in which *ToxhAT* was embedded. TSDs bounding *ToxhAT* were not identified in any of the three *B. sorokiniana* isolates, which left the question of whether *ToxhAT* remains an active transposon unanswered (20).

Whilst each of the three species that currently harbour *ToxA* have additional mechanisms of pathogenicity, the acquisition of *ToxA* is hypothesised to have enabled the emergence of each species as a significant pathogen of wheat. As such, understanding the mechanisms by which *ToxA* is horizontally transferred and/or remains in an active transposon is of great importance to future management of these diseases. In this work we sequenced and additional eight *B. sorokiniana* isolates and compared these with *P. tritici-repentis* genomes to understand the mobility of *ToxhAT* in these two species and explore the hypothesis that *ToxhAT* is an active class II transposon in *B. sorokiniana*.

## Results

### Chromosome-level assembly of eight *B. sorokiniana* isolates

Eight novel *ToxA^+^ B. sorokiniana* genomes were sequenced with the Oxford Nanopore MinION sequencer. The publicly available, fully assembled and annotated genome for the Australian *B. sorokiniana* isolate CS10 [BRIP10943, SAMN05928353] was used as a high-quality reference genome and the -*ToxA* isolate CS27 [BRIP27492, SAMN05928351] was also included for comparative analyses (20). The completeness of each new isolate was scored using the Dothideomycetes Benchmarking Universal Single-Copy Orthologues (BUSCOs) containing 3786 conserved orthologs (23). QUAST was used to calculate the fraction of CS10 genome captured in each isolate, giving an isolate-specific completeness score (24). The BUSCO and QUAST scores of the novel isolates fell within a narrow range, with a minimum of 93.8% BUSCO score and a minimum QUAST score of 93.5%. A summary of genome assembly completeness for each isolate is outlined in Table 1.

**Table 1:** Summary of Isolates collection details and genome assembly statistics.

| Isolate | Collection Year | Collection Location | Raw Data (Gb) | Genome Size (Mb) | Complete Chromosomes | BUSCO Score (%) | Quast Genome fraction (%) |
| --- | --- | --- | --- | --- | --- | --- | --- |
| CS10 (BRIP10943)* | 1966 | Biloela, QLD, Australia | NA | 36.92 | 16 | 99.6 | N/A |
| WAI2411 | 2015 | Nyngan, NSW, Australia | 14.20 | 36.02 | 10 | 95 | 94.911 |
| WAI2431 | 2015 | Gilgandra, NSW, Australia | 3.13 | 36.10 | 8 | 94.5 | 95.202 |
| WAI2432 | 2015 | North Star, NSW, Australia | 10.46 | 36.47 | 8 | 94.2 | 93.252 |
| WAI3285 | 2017 | Inglestone, QLD, Australia | 2.99 | 35.88 | 6 | 93.8 | 95.086 |
| WAI3295 | 2017 | Moonie, QLD, Australia | 2.84 | 36.02 | 4 | 94.5 | 94.263 |
| WAI3382 | 2016 | Pallamallawa, NSW, Australia | 10.09 | 36.21 | 9 | 95.7 | 94.966 |
| WAI3384 | 2016 | Mungindi, QLD, Australia | 12.67 | 36.76 | 12 | 95.4 | 93.603 |
| WAI3398 | 2016 | Walgett, NSW, Australia | 10.96 | 36.27 | 10 | 95.6 | 93.625 |

\* CS10 is an abbreviated name for isolate BRIP10943, (*Cochliobolus sativus* isolate 10)

Chromosome numbers were assigned to contigs in the new assemblies based on the numbering and orientation of the CS10 reference genome. Whole genome alignments identified chromosomes (chr) 1–16 and contig (tig) 17 in all new assemblies (Table 1, Supplemental Dataset 1). In six of the eight Nanopore isolates chr12 (1.85 Mbp) and tig17 (819 kbp) were present as a single contig totalling 2.5–2.7 Mbp. In CS10, chr12 contained two ribosomal repeats at one end and similarly tig17 also contained two ribosomal DNA repeats. We confirmed these two contigs were connected by the rDNA region by aligning the raw reads back to the assembled contigs to confirm the Nanopore assemblies manually (Figure S1A). Using this data chr12 and tig17 were taken to be one chromosome and were concatenated in all final Nanopore assemblies, labelled as chr12_tig17. Most isolates showed a high degree of synteny to the CS10 assembly, however WAI2432 contained several large chromosomal rearrangements. These were investigated and confirmed by aligning raw nanopore reads to the *de novo* assembly (Figure S1B).

### Re-sequencing of *B. sorokiniana* revealed a conserved sub-telomeric region

Chromosome completeness was assessed based on the presence of 5’ and 3’ terminal telomeric repeats. *B. sorokiniana* has a 6 bp telomeric repeat of “AACCCT” extending up to 450 bp, as identified in the CS10 reference genome (20). Whole chromosome alignments of the nine assemblies revealed a conserved sub-telomeric sequence. The region ranged from 500 bp in chr05 to >1 kbp in all other chromosomes and had a pairwise sequence identity of 88.2% (Figure S2). Chr08, chr09, and chr12 showed evidence of RIP mutation across the 3’ end of the telomeric region (see 380–1000 bp in Figure S2). Excluding these chromosomes from the analysis, the sequence was 96.5% identical across the remaining 11 chromosomes. The sub-telomeric region was identified at multiple chromosome termini of the newly sequenced isolates with >90% pairwise identity. This sub-telomeric sequence appears to be unique to *B. sorokiniana* as it was not identified in the publicly available *P. nodorum* isolate SN15 nor the *P. tritici-repentis* isolate 1C-BFP, two close relatives of *B. sorokiniana*. Similarly, a BLASTn analysis in the NCBI nr database found no significant hits outside of the *B. sorokiniana* genome. As this region was consistently identified adjacent to the telomeres in nearly every chromosome of CS10, it was used as a second marker of chromosome completeness for our nanopore assemblies. We considered nanopore assembled chromosomes complete if both termini had a minimum of three telomeric repeats (visually identified) or if ≥650 bp of the sub-telomeric region was present at the termini with ≥90% identity. Genome assembly size and completeness for each isolate is summarised in Table 1.

### *ToxhAT* is an active transposon within *B. sorokiniana*

*ToxhAT* was manually annotated in each genome using the CS10 annotation as a reference. The full transposon was present as a single copy in all newly sequenced isolates, with pairwise nucleotide identities of ≥99.5%, when aligned to CS10. All manually annotated genes within *ToxhAT* were present, with pairwise nucleotide identity of ≥98%. Two *ToxA* haplotypes were identified in the newly sequenced isolates, both matched previously documented *ToxA* sequences (16, 25). *ToxhAT* was identified in a different genomic location in all re-sequenced *B. sorokiniana* isolates, mostly in different chromosomes (Figure 1A). Unexpectedly, whole chromosome alignments also identified a conserved ∼200 kbp region carrying *ToxhAT* in all nine isolates (Figure 1A). We confirmed chromosomal location of the 200 kbp region in each chromosome, including *ToxhAT*, through the alignment of the raw reads to each *de novo* assembly (Figure S3).

**Figure 1.**
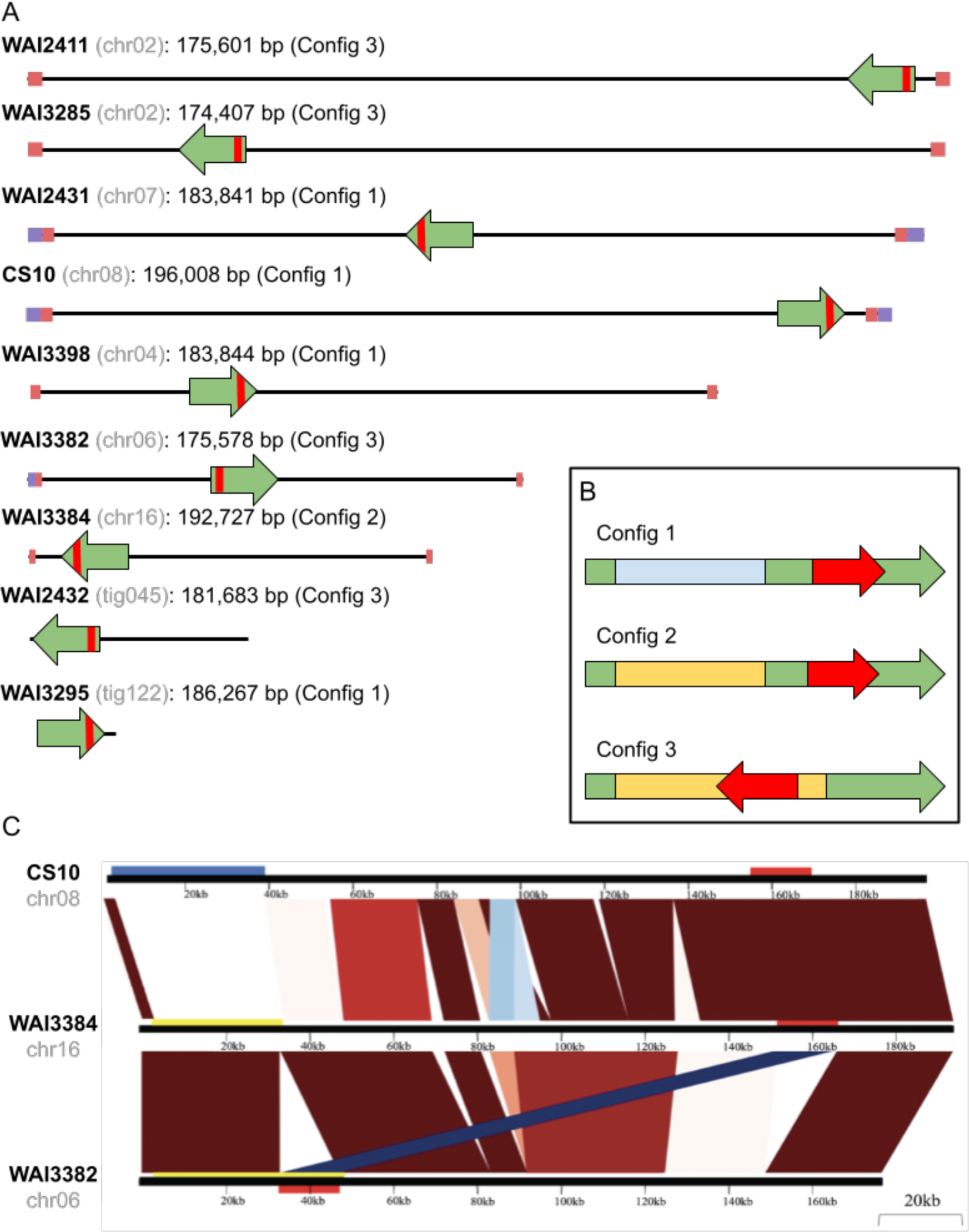
**A)** A schematic overview of the chromosomal location of *ToxhAT* and the conserved 200 kbp region that is shared between all re-sequenced *B. sorokiniana* isolates. The name of the isolate is shown in bold text with the chromosome or contig name in (grey). The exact size of the conserved element is shown in bp next to the chromosome name. Chromosomes containing telomeric repeats are indicated with a purple square, and the conserved sub-telomeric sequences are indicated with a pink square. The direction of insertion of the conserved 200kb is shown by the green arrow. The location of ToxhAT is shown by the small red square inside the green arrow. **B)** Insert: A schematic overview of the three “configurations” of the conserved mobile element. Sequence that shares strong homology >90% is indicated by the same color. ToxhAT is shown in red with its orientation indicated by the arrow. **C)** Whole chromosome alignment of each configuration showing highly-syntenic blocks in the same direction (red) and in the reverse direction (blue). The black horizontal bar denotes the chromosome. The horizontal blue bar in the top frame on CS10 chr08 shows sequence that is unique to config1 and the yellow bar in the bottom two frames shows sequence that is unique to configs 2 and 3. The location of *ToxhAT* is shown with the horizontal red bar.

To investigate how this 200 kbp may have translocated or moved between different chromosomes, this region was extracted from each isolate and realigned with all sequences in the 5’ to 3’ orientation as it is found in CS10 chr08. This alignment showed that these 200 kbp blocks had conserved edges. The exact length of the conserved DNA was 174–196 kbp and existed in three distinct configurations (configs) (Figure 1B, Table 2). Configs 1 and 2 were defined by the presence of *ToxhAT* at a shared 3’ position, whereas in config3 *ToxhAT* was inverted and translocated to a 5’ position within the larger 200 kbp. Config 1 could be distinguished from the other two by a unique 35 kbp sequence that was not present in the other two configs. In configs 2 and 3 this sequence was replaced by a different 30 kbp sequence (Figure 1B-C). Excluding these regions which were unique to either config 1 or configs 2 and 3, the remaining 160 kbp DNA alignment showed ∼ 91.3% pairwise nucleotide identity.

**Table 2.** Chromosomal location, length, orientation and a summary of gene content in each config.

| Isolate | Chromosome | Min (bp)<sup>1</sup> | Max (bp)<sup>2</sup> | Length (bp) | Orientation<sup>3</sup> | *ToxhAT* Location<sup>4</sup> | Total genes annotated | Config | Av. config length (bp) | Av. Gene count per config |
| --- | --- | --- | --- | --- | --- | --- | --- | --- | --- | --- |
| CS10 | chr08 | 1875320 | 2071325 | 196005 | + | 3' | 55 | Config 1 | 187479 | 52 |
| WAI2431 | chr07 | 1731625 | 1915465 | 183840 | - | 3' | 52 |  |  |  |
| WAI3295 | tig122 | 1 | 186228 | 186227 | + | 3' | 49 |  |  |  |
| WAI3398 | chr04 | 633088 | 816931 | 183843 | + | 3' | 51 |  |  |  |
| WAI3384 | chr16 | 165966 | 358692 | 192726 | - | 3' | 57 | Config 2 | 192726 | 57 |
| WAI2411 | chr02 | 3435149 | 3610749 | 175600 | - | 5' | 52 | Config 3 | 176816 | 53 |
| WAI2432 | tig045 | 42029 | 223711 | 181682 | - | 5' | 54 |  |  |  |
| WAI3382 | chr06 | 734719 | 910295 | 175576 | + | 5' | 52 |  |  |  |
| WAI3285 | chr02 | 1563991 | 1738397 | 174406 | + | 5' | 52 |  |  |  |

<sup>1</sup> Chromosomal position that is the start of the config
<sup>2</sup> Chromosomal position that is the end of the config
<sup>3</sup> With reference to orientation of CS10 chromosomes, whether the conserved start of the config is on the positive or negative strand
<sup>4</sup> Location of ToxhAT within the config

As *ToxhAT* was present in two distinct locations within the larger conserved DNA element, we investigated the empty insertion sites for evidence of active translocation via a transposase, namely target-site duplications (TSDs). A multiple sequence alignment of all 9 *ToxhAT* sequences from configs 1 & 2 (3’ position) against config 3 (5’ position) identified a 2 bp TSD of “TA” bounding the transposon (Figure 2A). In contrast the empty insertion sites had only one “TA” present (Figure 2B). This shows that *ToxhAT* is likely an active transposon with a two-base pair “TA” TSD.

**Figure 2.**
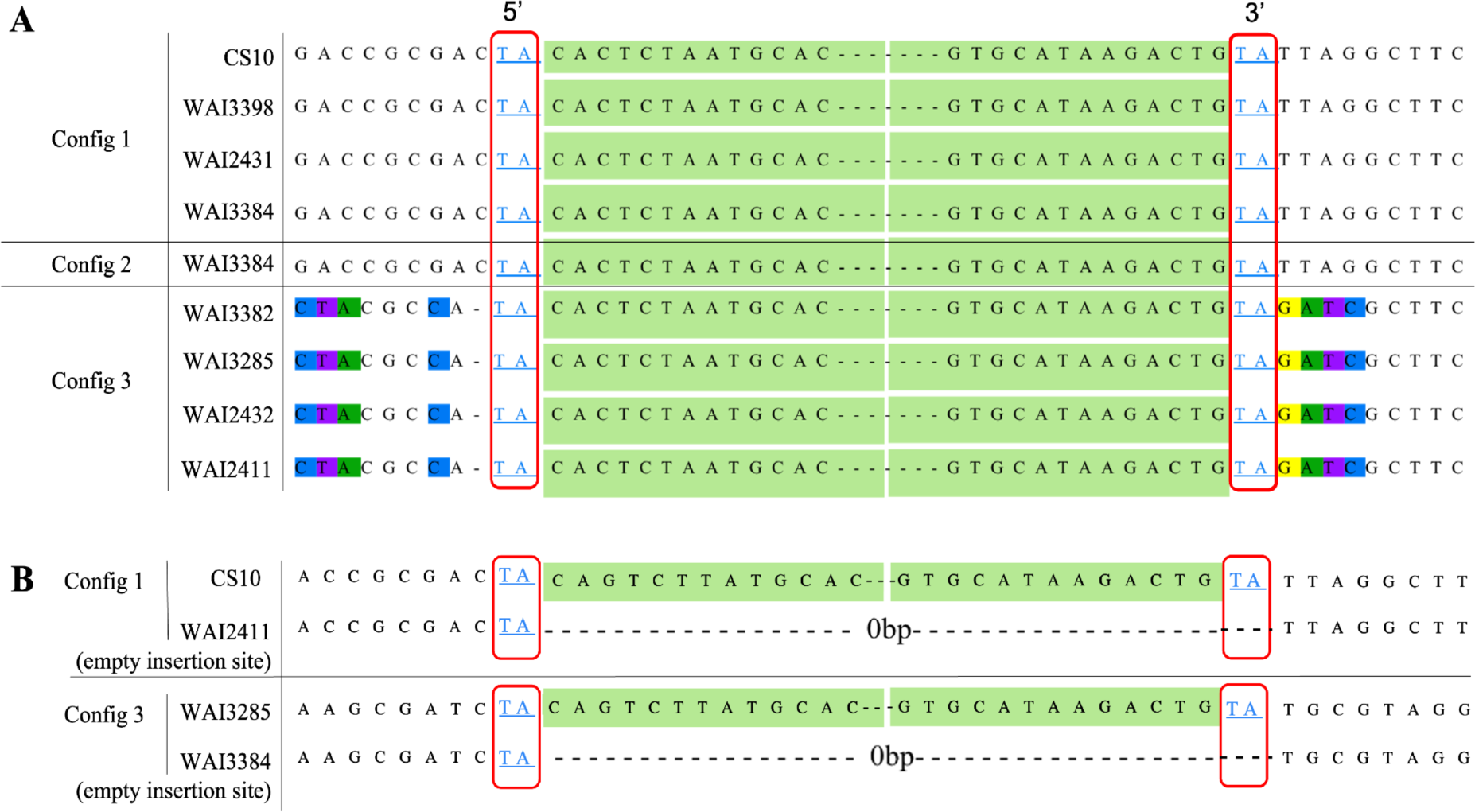
**A)** Alignment of all re-sequenced isolates showing a partial sequence of the TIR of ToxhAT in green. The “TA” target site duplication found on either side of ToxhAT in all three configurations is highlighted in the red box. **B)** Alignment of the ToxhAT to an empty insertion site in a different isolate. The “TA” TSD only appears once in the empty insertion site, but twice on either side of *ToxhAT,*.

### *ToxhAT* is moving as cargo within a *Starship* transposon

After investigating the movement of *ToxhAT* alone, we next sought to investigate the putative mechanism driving the movement of the larger conserved 200 kbp element. Large genomic rearrangements in fungi can be attributed to many different mechanisms, including but not limited to ectopic recombination, unequal crossing over, breakage-fusion-bridge cycles and transposon-mediated transposition (26). Given the high frequency of movement and conserved boundaries, two features often associated with transposons, we explored the hypothesis that these chromosomal rearrangements were driven by transposons (9, 13, 27). First, all genes identified within the 174–196 kbp of all three configs were manually annotated and assigned a putative function based on conserved domains (Supplementary Dataset 2). Excluding genes within *ToxhAT*, 59 unique genes were identified across all three configurations (Table 2).

Notably, the first gene at the 5’-end in all three configs was predicted to encode a DUF3435 domain. Genes that encode this domain have been proposed by Gluck-Thaler *et al.* (28) to be a novel group of transposases that mobilise large transposons now called “*Starships*” (8, 28–30). *Starships* are defined as large mobile elements, whose conserved DUF3435 gene, referred to as the “Captain” is the first gene encoded at the 5’ edge of the transposon. In all three configs, the DUF3435 Captain is found ∼350 bp from the 5’ edge and the average pairwise identity of the gene was 96% across all nine isolates. The gene product is predicted to contain a YR domain at the middle of the translated product and a C-terminal DNA-binding domain (Figure S4). In addition to the YR-captain, *Starships* also contain several other conserved protein families. These include ferric-reductases (FREs), NOD-like receptors (NLRs), patatin-like phosphatases (PLPs) and DUF3723 proteins (28). These “accessory” genes in *Starship* transposons were also present in all three configs, further supporting the classification of this mobile region as a novel *Starship*. This included three PLPs and three predicted NLRs (Supplementary Dataset 2).

*Starship* transposition has now been shown to be mediated by the YR-encoding Captain (8). This mechanism results in the formation of a TSD variant known as short direct repeats (SDRs). Both TSDs and SDRs are 4–6 bp in length and flank the ends of the inserted transposon. However, during excision while both TSDs are left at the excision site, only one SDR remains as the other is excised with the transposon. As such, the conserved boundaries of *Sanctuary* were analysed for the presence of SDRs. Close examination of the sequence immediately upstream of the 5’-end revealed a 6 bp sequence which was present in the empty insertion sites of all isolates (Figure 3A). This 6 bp sequence was not perfectly conserved but had a consensus sequence of “ATWHCT”, where W is an A or a T and H is an A, T or C. All nine re-sequenced isolates showed perfect conservation of the last six base pairs, which we now refer to as the 3’ SDR (“ATACCT”). Based on the high degree of conservation at the 3’ end we hypothesise that *Sanctuary* excises through a recombination driven mechanism, carrying the 3’ SDR sequence with the transposon as it moves (drawn here as an extrachromosomal circular DNA) (Figure 3B). Attempts to PCR a circular intermediate sequence from CS10 grown under different environmental conditions were not successful in obtaining a specific amplicon (data not shown). Further optimisation of the conditions required to induce *Starship* movement is required before this hypothesis can be explored in more detail.

**Figure 3.**
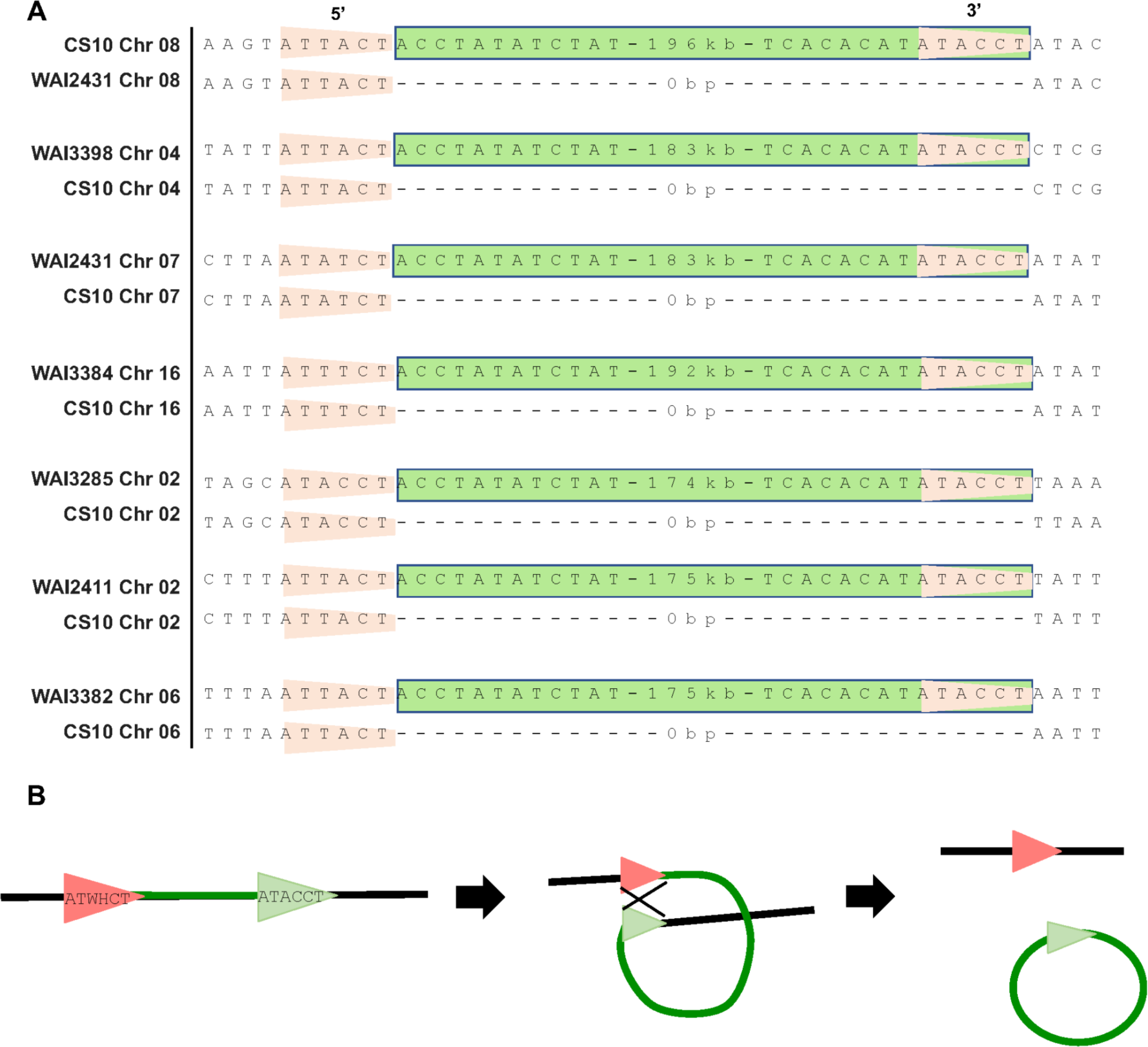
**A)** Alignment of the edges of *Sanctuary* to an “empty” site from the same location in another isolate. Isolate names and chromosomes are given on the far left, the 6- bp SDR is shown in the orange triangle and the truncated *Starship* sequence is highlighted in the green box with its size in bp shown. Empty insertion sites are shown with dashes to align the edges. **B)** Schematic of hypothesised recombination driven excision from the chromosome. The 5’ target site (orange arrow) remains in the chromosome after excision, while the 3’ SDR (green arrow) moves with the transposon.

### Mini-*Sanctuary* transposons are present in *B. sorokiniana*

Using the Captain gene as a search query, we identified two “mini” *Sanctuary* transposons (mini-*S1*). A conserved 72.7 kbp mini-*S1* was found at the same chr 03 locus in all nine *ToxA B*. *sorokiniana* genomes. A 47.5 kbp mini-*S1* was identified on chr 08 in *toxa-* isolate CS27. The latter was at a similar, but not identical location, as the full-length *Sanctuary* in CS10 (Figure 4A). Alignment of the mini-*S1s* with the full-length *Sanctuary* found that the first 3.8 kbp, including the Captain, was conserved with a sequence identity of 98.5%. The final 3.1 kbp was conserved with a sequence identity of 88.7%, including *S1-50*, a predicted chromodomain-containing protein. Annotation transfer also found multiple transposons embedded within full-length *Sanctuary* were also present in the mini-*S1s* (Figure 4A, Table S1). Outside of these conserved regions and homologous genes, the mini*-S1s* shared little sequence similarity with each other and with *Sanctuary* (<70%). However, whilst the first four bases of both mini-*S1s* differed slightly (“AGTA” in mini-*S1s* and “ACCT” in *Sanctuary*) the 5’- and 3’-SDRs were conserved at both termini (Figure 4B and C). Given the conserved nature of the boundaries, the Captain, and *S1-50*, these components may constitute the minimal unit required for transposition. Mini-S1s were not identified in any species outside of *B. sorokiniana* via NCBI BLASTn search.

**Figure 4.**
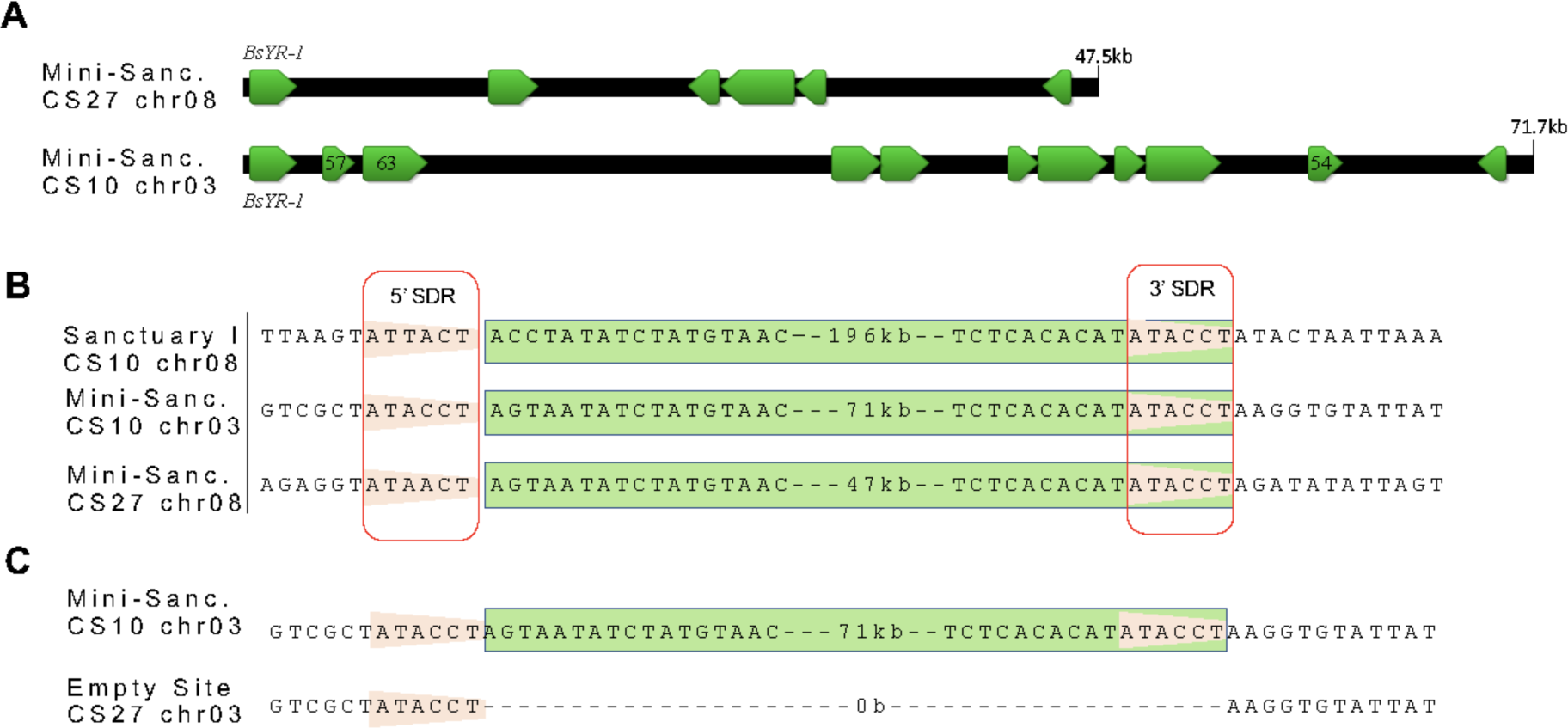
**A)** Schematic overview of Mini-S1 found in the *toxA-* isolate CS27 and the conserved mini-S1 found in chr 03 of all *ToxA+* isolates. Annotated genes are shown in green. **B)** Alignment of the edges of *Sanctuary* I from CS10 to both mini-S1 *Starship*s shown in part A. The 6 -bp SDR consensus sequence remains the same at the 5’ end of each element and the 3’ SDR is perfectly conserved. **C)** Comparison of the empty site in CS27 at Chr. 3 to the insertion of mini-S1 at the same location in CS10.

### Structural predictions of the *Starship* YRs show similarity to the bacteriophage P1 Cre recombinase

Our discovery of *Sanctuary* in *B. sorokiniana* occurred concurrently with the discovery of a second *Starship*, *Horizon,* that was found to harbour *ToxhAT* in *Pyrenophora tritici-repentis* (31). Attempts to align the nucleotide sequence of *Sanctuary* and *Horizon* revealed no significant similarity, except for the presence of *ToxhAT.* We next sought to look for homology at the amino acid level for each Captain and again detected little homology between the two proteins. In the absence of strong homology at the sequence level, we next sought similarity using structural prediction. The 786 aa Captain from *Sanctuary* was predicted by NCBI BLASTp and Phyre2 protein fold recognition server to contain a site-specific YR domain from 111– 463 with 98.8% confidence, and a DNA-binding domain from 711–784 with 95% confidence (Figure S4). In order to explore the DNA-binding capacity of these putative transposases in more detail, structural predictions of both the *Sanctuary* and *Horizon* YRs were generated using AlphaFold2 (Figure S5) (32). To identify whether our predicted YR models shared any structural similarity to any existing structures in the Protein Data Bank (PDB), we submitted these models to the DALI protein structure comparison web server (33). Structural similarity was assessed based on the root mean square deviation (RMSD) and number of equivalent residues (LALI). Using these metrics, both *Starship* YR models were found to share structural similarity with the crystal structure of the bacteriophage P1 Cre recombinase bound to loxP DNA (34). The DALI server results for *Sanctuary* included the P1 Cre recombinase (PDB: 3C29) with a RMSD of 3.5 Å and LALI of 264, indicative of a shared structural fold. *Horizon* also showed structural similarity to the P1 Cre recombinase (PDB: 3MGV) with a RMSD value of 3.3 Å and LALI of 221. While some regions of both YR monomer models were well predicted, the active site of the *Sanctuary* YR was not. Previous structural analyses of the P1 Cre recombinase demonstrates this YR to assemble into a tetrameric complex (34). To enhance the confidence of the active sites in both *Starship* YRs, AlphaFold2 multimer (35) was employed to generate tetrameric models for *Sanctuary* (Figure 5A) and *Horizon* (Figure 5B). These YR tetramer predictions yielded models with higher global confidence compared to monomer models and improved model confidence around the putative active sites. Furthermore, superimposition of the *Starship* YR tetrameric models (Figure 5C) and the P1 Cre recombinase crystal structure (Figures 5D and S6A) demonstrates the similarity in stoichiometry and domain arrangement between these YRs, despite the *Starship* YRs being larger.

**Figure 5:**
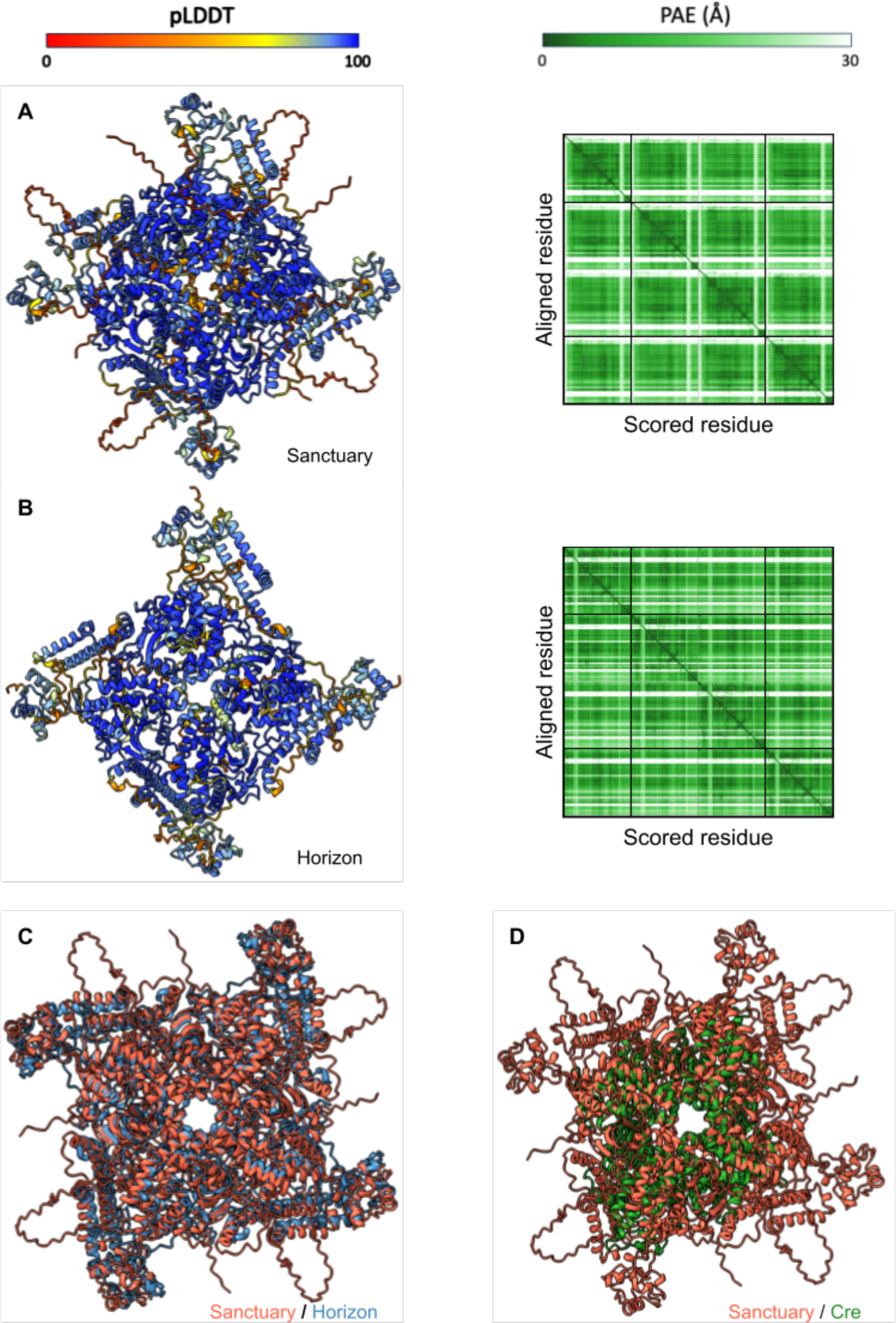
Tetrameric AlphaFold2 predictions of the *Sanctuary* and *Horizon* tyrosine recombinases demonstrate structural homology to Cre recombinase. AlphaFold2 predicted tetrameric models of *Sanctuary* (A) and *Horizon* (B). The corresponding predicted aligned error (PAE) heatmap is shown to the right of both models, measured in Ångstrom (Å). Models are coloured by confidence, represented by predicted local distance difference test (pLDDT). Superimposition comparison between *Sanctuary* and *Horizon* YRs (C) and *Sanctuary* and P1 Cre recombinase (PDB: 3MGV) (D) are also included.

The P1 Cre recombinase (Figure 6A) uses a large basic surface to bind DNA (Figure 6C), with the active residues (R173, E176, K201, H289, R292, W315, Y324) residing at the interface with the DNA molecule (Figure 6B) (34). Analysis of the superimposed models of the *Sanctuary* tetramer prediction and the structure of the P1 Cre recombinase bound to loxP DNA indicates the same basic surface used to bind DNA exists in the fungal YR (Figure 6E). While the structural prediction of the fungal YRs cannot be performed to incorporate DNA models, comparison with the Cre recombinase structure suggests the R-Y-R-Y (R260, Y424, R427, Y457) tetrad of *Sanctuary* is also located in proximity to the DNA binding interface, as expected (Figure 6D). Further investigation into the predicted *Sanctuary*-DNA interface has revealed possible other residues-of-interest (E45, S171, K308, K456), which may contribute to the DNA-binding interface (Figure 6D). We could not identify the *Horizon* R-Y-R-Y tetrad by sequence analyses, however the structural alignment of the P1 Cre recombinase active site and the corresponding region in *Horizon* reveals a putative R-Y-R-Y tetrad of residues R84, Y112, R292, and Y322 (Figure S6). The combined analyses of the predicted structures of both *Starship* YRs confirm the existence of a YR fold, even in the absence of DNA sequence similarity, and strongly indicate the presence of functional DNA binding interfaces.

**Figure 6:**
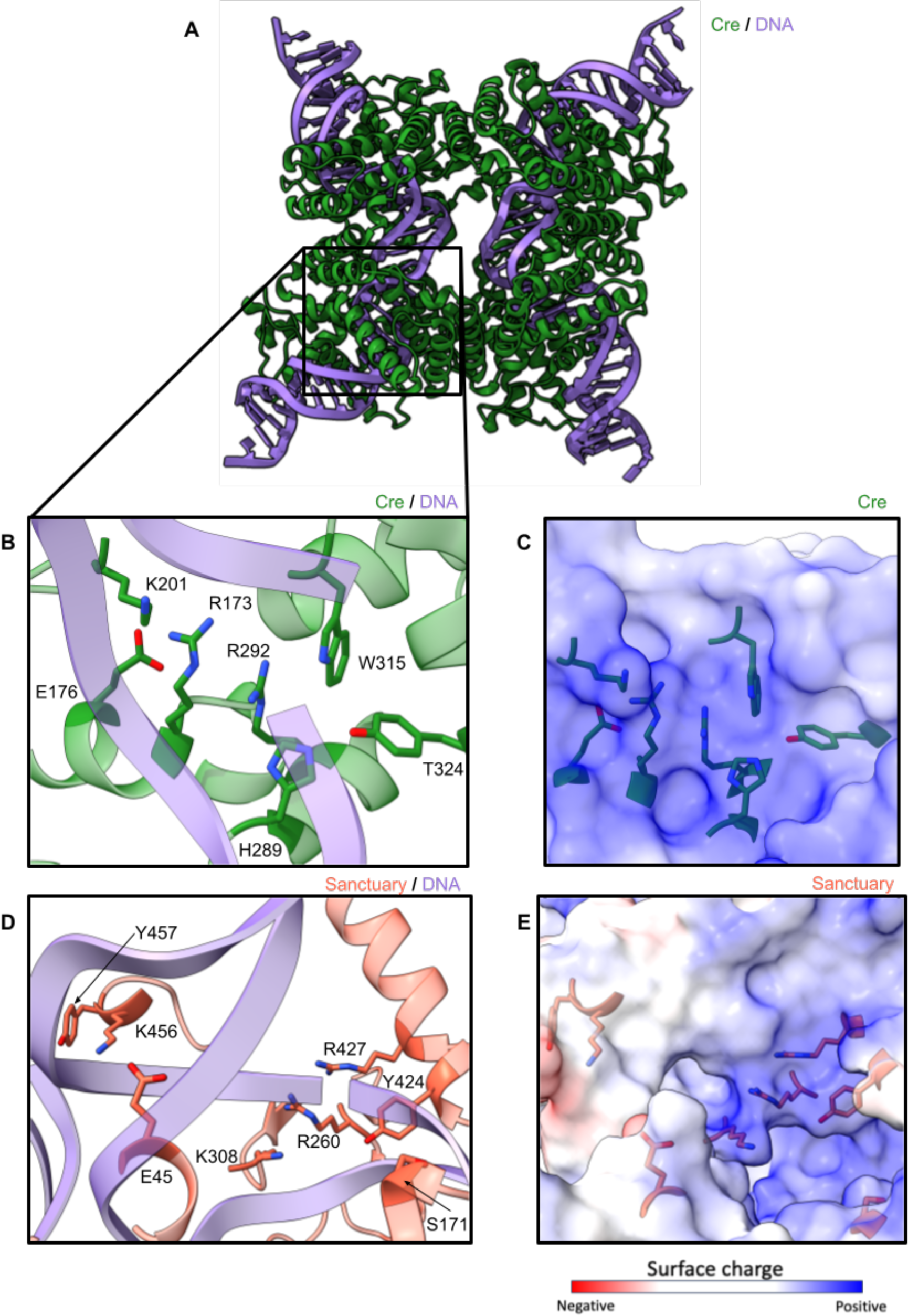
The *Sanctuary* AlphaFold2 tetrameric model maintains a DNA binding site comparable to P1 Cre recombinase. (A) Ribbon structure of the P1 Cre recombinase (PDB: 3MGV) including DNA interfaces. (B) Zoom in on the DNA (purple) binding interface of the P1 Cre recombinase (green) including highlighted catalytic residues. (C) Surface structure coloured by surface charge of the P1 Cre recombinase DNA interface showing catalytic residues (green). (D) Predicted *Sanctuary* YR DNA binding interface (orange) with superimposed DNA from the P1 Cre recombinase model (purple). Previously identified catalytic residues from sequence data are highlighted alongside putative residues-of-interest identified from structural analysis. (E) Surface structure coloured by surface charge of the predicted *Sanctuary* model interface showing catalytic residues (orange).

## Discussion

This study presents new insights into the previously discovered class II *ToxhAT* transposon that carries the agronomically-significant *ToxA* effector. Here, we found evidence that *ToxhAT* is an active transposon within *B. sorokiniana* based on the identification of a two bp “TA” TSD. In addition, we have found that in *B. sorokiniana ToxhAT* is ‘cargo’ within a much larger *Starship* transposon, now named *Sanctuary*. There are multiple versions of *Sanctuary* found within the *B. sorokiniana* genomes sequenced thus far, including two “mini” *Sanctuaries* present in both *ToxA+* and *toxa-* isolates. A comparative analysis of the larger HGT event between all three fungal species known to carry *ToxhAT,* shows that this transposon has been captured independently by two distantly related *Starship* transposons, *Sanctuary* in *B. sorokiniana* and *Horizon* in *P. tritici-repentis* and *P. nodorum* (31).

With only one TSD present within empty insertion sites, but two flanking *ToxhAT* sequences, we conclude that *ToxhAT* is mobile within all sequenced *B. sorokiniana* isolates. However, the discovery of the “TA” TSD sequence brings the classification of *ToxhAT* into question. A *hAT*-like transposase was reported when *ToxhAT* was initially discovered in *P. nodorum* (15). Typically, *hAT*-type transposons have 8 bp TSD and TIR sequences that range in size from 5-27 bp (27). This contrasts with *ToxhAT* having the 2 bp TSD discovered in this study, and 74 bp TIR sequences reported previously (20). These characteristics could be explained either by *ToxhAT* being a non-canonical *hAT* transposon, or that it’s been misclassified. The 2 bp TSD and 74 bp TIR of *ToxhAT* is representative of the *Tc1/Mariner* type of class II transposons. The *Tc1/Mariner* superfamily is characterised by having the same 2 bp “TA” TSD (36) as *ToxhAT*. While typical *Tc1/Mariner* transposons have short TIRs between 20–30 bp (37), some have been identified with TIRs up to 169 bp (38). Despite many attempts to manually annotate a transposase within *ToxhAT,* we have been unable to identify any predicted gene that strongly matches a transposase in either the *hAT* or *Tc1/Mariner* superfamilies. Therefore it is difficult to confidently classify *ToxhAT* as belonging to either group based on the TSD alone.

In all nine *ToxhAT*-positive *B. sorokiniaina* isolates, *ToxhAT* was embedded within a conserved ∼200 kbp region. The region, designated *Sanctuary*, is a *Starship* transposon. *Starships* are a novel group of transposons characterised by their large size, a DUF3435-containing YR as the first gene in the 5’ direction, and for having a cargo and accessory gene structure (28). Cargo genes vary between elements but generally confer fitness advantages to the fungal host in specific environments, while accessory genes fall into several conserved gene families all of which are found within *Sanctuary* (28, 30). *Sanctuary* was found in three separate configurations within the nine isolates, but all configurations contain the same “ATWHCT” SDR sequence suggesting it is one element. Surprisingly *Sanctuary* is the second identified *Starship* carrying *ToxhAT*, the first being *Horizon* in *P. tritici-repentis* and *P. nodorum* (31). While both *Sanctuary* and *Horizon* carry *ToxhAT*, outside of this region there is no sequence similarity between them. This indicates that *ToxhAT* has been captured twice, independently, by *Horizon* and *Sanctuary*. The process that drives the ‘capture’ of cargo genes, such as *ToxhAT,* into *Starships* remains uncharacterised. The presence of this important virulence factor in two different *Starships,* adds to the growing body of evidence that *Starships* are facilitating HGT between fungi, and that the cargo that they carry confers a fitness benefit to the fungal host (8, 28).

Herein, we also show that in *B. sorokiniana* there are naturally occurring mini-*Sanctuary* elements. These mini elements are considerably smaller than the *Sanctuary* elements that carry *ToxhAT*, and do not contain most cargo and accessory genes. The mini-*Sanctuary* elements do have the same six base-pair SDR as the larger element, which also supports their classification within the same family (39). It remains unclear at this stage if these mini-*Sanctuary’s* also remain active transposons as in the isolates sequenced for this study these elements were found in a conserved genomic location in all *ToxA* containing isolates. Urquart *et al.* (8) recently engineered a mini-*Starship* containing only the Captain transposase and the edges of the transposon and showed that this minimal unit is mobile in its host genome. Therefore, we hypothesise that our naturally occurring mini-*Starship*s might also remain active transposons. Their presence in *B. sorokiniana* gives us an opportunity to track the evolutionary history of *Sanctuary* at the population level and potentially reveal further sequence intermediates that more closely resemble the full *ToxhAT* carrying *Sanctuary.* These population level studies could provide important clues as to how *Starships* gain and lose genes in their natural fungal host.

The structural modelling of *Sanctuary* and *Horizon* has provided new and important insights into how the site directed recombination reaction may occur based on the strong structural similarity to the P1 Cre recombinase. It has already been proposed that *Starships* move via a site-directed recombination event between the short-direct repeats that form the boundaries of most *Starships* described thus far. Several groups have identified core catalytic residues that are found in other YR-domain containing enzymes, including Cre, but also Crypton transposases (8, 28, 40). These site-directed recombinase enzymes bind SDRs to facilitate a recombination event that results in one circular molecule containing one copy of the SDR, while the remaining SDR stays in the linear chromosome. Urquart *et al.* (8) have already shown that mobilised mini-*Hephaestus* elements do leave behind a “clean excision” after moving to a new genomic location and we have postulated that the *Starships* exist as extra-chromosomal circular DNAs (eccDNAs) after excision from the genome (41). Our AlphaFold modelling of both *Sanctuary* and *Horizon* Captains shows strong structural similarity to the site-specific P1 Cre recombinase (42). P1 Cre recombinase binds a conserved 34 bp *loxP* site and is one of the best studied site-specific recombinases (34). Using AlphaFold Multimer, we obtained a high-confidence model that suggests that both *Sanctuary* and *Horizon* Captains, like Cre, bind their DNA targets as a tetramer. These models enabled us to rapidly identify putative DNA binding residues that match the described Y-R-Y-R. These results highlight the growing power of structural bioinformatics to assist in the identification of key catalytic residues without the need to perform large scale alignments, which are technically difficult with more distantly related proteins.

Unlike other transposons, it has been argued that horizontal movement of *Starships* is required for them to persist over evolutionary time-scales (8). Under this hypothesis, cargo genes are essential for *Starships* survival and therefore confer fitness benefits to the fungal host when present. What remains less clear is the role and function of accessory genes within *Starships.* Examples of accessory genes include PLPs and NLRs, some of which have links to self/non-self recognition pathways in filamentous fungi and have been proposed to involved or required for horizontal movement (8). *Sanctuary* contains many accessory genes with these domains and is now one of a few characterised *Starships* where there is evidence of active transposition in natural isolates. This makes *Sanctuary* an exciting target for attempting to induce HGT events to better understand this process.

## Materials and Methods

### Fungal Culture and DNA extraction

*B. sorokiniana* isolates were obtained from NSW Department of Primary Industries, Wagga Wagga Agricultural Institute (WAI). These isolates and details pertaining to the year and site of collection are provided in Table 1. Fungal cultures were grown on V8-PDA media at 22°C under a 12 hr light/dark cycle. Cultures ranging in age from 5-10 days were scraped from the agar surface using a sterile razor blade. Harvested cultures were lyophilised for 48 hr and high molecular weight (HMW) DNA was extracted using a modified method from Fulten *et al.* (43) and our full protocol is available here: dx.doi.org/10.17504/protocols.io.k6qczdw.

### Genome sequencing, *de novo* assembly and scaffolding

Genome assemblies for isolates BRIP10943a (CS10) and BRIP27492a (CS27) were obtained from McDonald *et al.* (20). All new raw data generated for this study were deposited in NCBI BioProject ID PRJNA1017791. HMW DNA for each isolate was sequenced using the Oxford Nanopore MinIon R9.4 Flow cells with the 1D library kit SQK-LSK08 (44). All DNA samples were cleaned three times using Agencourt AMPure beads prior to starting the 1D library kit (Beckman Coulter, Inc., CA, USA). Genomes were assembled with Canu v1.6 with a minimum read length of 5 kbp (45). *De novo* genome assemblies were corrected using the trimmed reads output from Canu. Trimmed reads were mapped to the genome with Minimap2 v2.17 (46) followed by consensus calling with Racon (47). The output consensus sequence from Racon was used as input for iterative correction five times.

Contigs in the polished Nanopore assemblies were assigned chromosome numbers based on the numbering and orientation of the CS10 reference genome (20). The CS10 assembly has 16 complete chromosomes labelled by descending size from chromosome 1–16. Six partial contigs were also present, named by descending size, contigs 17–22 (20). LastZ v1.04.03 implemented in Geneious was used to align assembled scaffolds to the CS10 reference genome (48). A contig was assigned a chromosome number if >90% of its length aligned to a single CS10 chromosome. In instances where multiple smaller contigs aligned in tandem to form a single CS10 chromosome, the contigs were scaffolded together. A summary of the number of contigs, genome size, presence of telomeres and final chromosome names for each isolate is provided in Supplementary Dataset 1.

### Assessing assembly accuracy and completeness

The quality of each polished assembly was assessed using BUSCO v 4.1.4, run with the Dothideomycetes dataset of (3786 genes) (23). BUSCO was run on the command line: busco -m genome -l dothideomycetes_odb10 --cpu 12 -i <isolate.fasta> -o <OUTPUT_DIRECTORY>. QUAST was used to compute the percentage of the CS10 genome assembly captured within each final Nanopore assembly (24). The online QUAST interface was used, accessible at http://cab.cc.spbu.ru/quast/. No contig size was excluded.

### Genome Annotation

*Funannotate* v1.7.4 was used to annotate whole genomes of all WAI isolates and the CS10 genome (https://funannotate.readthedocs.io/en/latest/). Briefly, all assemblies were cleaned using the *funannotate sort* and *funannotate mask* commands. The *funannotate train* command was then used to train the pipeline for each isolate using the cleaned assembly file and CS10 RNA-seq data (20). Finally, the *funannotate predict* command was used to annotate each isolate assembly using the trained pipelines. Max intron was set at 1000 bp for both train and predict and minimum gene size was kept defaulting 50 bp. The output of the seven individual gene programs used during the *funannotate* process are combined into one consensus file with Evidence Modeler (EVM). *Funannotate* uses “weighting” scores to instruct EVM on which output to accept when two software have conflicting annotations (49). The final weightings were kept to default except CodingQuary (default 2, adjusted 6), and Program to Assemble Spliced Alignments (PASA) (default 6, adjusted 4) (49). No iteration of the *funannotate* pipeline was able to annotate the genes encompassed by the *ToxhAT* transposon. *ToxhAT* was therefore manually annotated in each genome using the CS10 annotation as a reference.

### Putative gene function identification

Putative gene functions were identified using the ExPASy translation tool (https://web.expasy.org/translate/) and analysed for conserved domains using the online NCBI Conserved Domain Search tool (CDS, https://www.ncbi.nlm.nih.gov/Structure/cdd/wrpsb.cgi), and Phyre2 v2.0 (50). All searches used the Conserved Domain Database, a superset containing NCBI-curated domains, and the Pfam, SMART, COG, PRK, and TIGRFAM databases (51). The e-value threshold was set at max 0.01. Phyre2 was run in normal mode and hits with below 90% confidence in the model were excluded from analysis. Cut-offs for amino acid percentage identity were considered on a case-by-case basis. Hypothetical gene functions were assigned if a hit with a low e-value was provided by at least one of these three protein predictors.

### AlphaFold predictions and structural analyses

Monomeric predictions of the *Sanctuary* and *Horizon* tyrosine recombinases were performed using the AlphaFold2 ColabFold v1.5.5 (52), while tetrameric predictions were generated using AlphaFold2 Multimer (35), via a local installation of AlphaFold2 v2.3.2. The DALI protein structure comparison server was used to compare AlphaFold2 models to existing structures in the Protein Data Bank (33). Structural analyses were performed and visualised with ChimeraX (53).

### Data Availability

All data are available under NCBI projects number PRJNA505097 and PRJNA1017791. New BioSamples created for this manuscript are SAMN39994716-SAMN39994722. New Nanopore raw sequence data is in NCBI’s SRA Accessions SRR28048243-SRR28048237. Full genome assemblies including unplaced contigs and the mitochondrial genome are available at https://github.com/megancamilla/Sanctuary_manuscript.

## Acknowledgements

We thank The Sainsbury Laboratory for the use of their computational resources for generating AlphaFold2 models. The Sainsbury Laboratory is supported by the Gatsby Charitable Foundation, the University of East Anglia, and a BBSRC ISP grant in Advancing Plant Health (BB/Y002997/1). M. McDonald would like to acknowledge the Sun Foundation’s Peer Prize for Women in Science. K.C. Goncalves dos Santos acknowledges scholarships from the Fonds de Recherche du Quebec Nature et Technologie and, MITACS for her doctorate scholarships and, Centre SEVE and the UQTR for her internship scholarships. H. Germain was supported by Tier II Canada Research Chair CRC-2017-103.

